# TBpop: an open-access portal integrating genomic variation, population genetic statistics, phylogeny, pangenome composition, and strain metadata of *Mycobacterium tuberculosis* isolates from a population-representative survey in China

**DOI:** 10.64898/2026.08.06.742672

**Authors:** Yang Zhou, Fei Huang, Yanlin Zhao

**Author notes:** Corresponding authors: Yang Zhou,; Yanlin Zhao.

## Abstract

Tuberculosis remains a major global public health threat. While whole-genome sequencing has transformed our understanding of the causative agent, *Mycobacterium tuberculosis* (MTB), existing genomic databases are highly fragmented and often underrepresent structural variations (SVs). Furthermore, critical population-genetic statistics are rarely integrated with phylogenetic and geographic context, forcing researchers to reconcile separate datasets manually. To address this gap, we developed TBpop (https://tbpop.chinacdc.cn), an open-access, integrated population genomics portal. TBpop is built from 420 clinical MTB isolates selected from the first national drug resistance baseline survey in China. The portal integrates isolate metadata, pangenome categories, SNPs, SVs, IS6110 insertion sites, strain phylogeny, and gene-level statistics, and provides three interactive explorer modules: the Population Explorer, the Statistics Explorer, and the Variation Explorer. Additionally, a User Analysis module allows researchers to run population genetic workflows on their own alignments. TBpop provides an integrated platform for exploring genome plasticity, signatures of positive selection, and conservation patterns of functionally important genes in MTB.

## Introduction

The World Health Organization estimated that 10.7 million people developed tuberculosis (TB) and 1.23 million people died from TB in 2024, making TB the world’s leading cause of death from a single infectious agent and a major contributor to antimicrobial-resistance-associated mortality [1]. China is one of the major high-burden countries, accounting for an estimated 6.5% of global TB incidence in 2024 and therefore represents an important setting for genomic resources that can support studies of the causative pathogen *Mycobacterium tuberculosis* (MTB) in population structure, transmission, drug resistance, and pathogen evolution.

Whole-genome sequencing has transformed the study of MTB by enabling high-resolution reconstruction of strain lineages, transmission clusters, resistance-associated mutations, and evolutionary histories [2]. Existing MTB genomic resources have made important contributions to the field. Resistance-oriented resources such as ReSeqTB primarily support interpretation of antimicrobial-resistance mutations or isolate-level resistance evidence [3]. CRyPTIC consortium has compiled the single largest dataset of global MTB epidemic strain with matched whole-genome sequences and quantitative minimum inhibitory concentration measurements [4]. General MTB variation resources such as GMTV and tbvar provide nucleotide-level variation catalogues [5,6], while annotation resources such as Mycobrowser and TB Genome Annotation Portal have supported genome browsing and functional interpretation [7,8] . However, these resources usually focus only on SNPs and small indels; genomic structural variations (SVs) are often underrepresented despite their relevance to gene disruption, genome plasticity, lineage differentiation, and potential instability of clinically or immunologically important loci. In addition, many analyses of MTB genomics remain distributed across static supplementary tables, specialized resistance catalogues, command-line workflows, or single-purpose databases. This fragmentation makes it difficult for non-specialist users to move from a population-level signal to the underlying variants and then to a phylogenetic interpretation of those variants.

Population-genetic statistics provide another underused entry point for exploring MTB genomic data. Measures such as nucleotide diversity π, Tajima’s D, homoplasy count, and Fst can highlight genes with unusual evolutionary patterns, including elevated diversity, repeated mutation, population differentiation, or signals affected by demographic history and selection [9–11]. These statistics are most useful when they are not treated as isolated endpoints, but when interpreted together with phylogeny, geography, drug-resistance phenotype, and population structure. Without a linked interface, this type of interpretation requires users to reconcile separate files and figures manually.

TBpop was developed to address this gap as an open-access population genomics portal for MTB. The current resource is built from 420 clinical MTB isolates from China, derived from the first national drug-resistance baseline survey and previously analysed in studies of MDR-TB evolution and the non-redundant MTB pangenome [12–16]. The portal integrates isolate metadata, lineage and population information, H37Rv-based gene annotations, pangenome categories, SNPs, SVs, IS6110 insertion sites, a strain phylogeny, and gene-level statistics. By organizing these data around linked genes, genomic coordinates, samples, and phylogenetic context, TBpop allows users to move from broad population patterns to specific genomic events without switching between independent datasets. Therefore, the main goal of TBpop is not to replace existing resistance knowledgebases or raw sequence repositories, but to provide a complementary exploratory resource for MTB population genomics. On TBpop, users can browse geographic and lineage structure, filter genes by pangenome category or evolutionary statistic, inspect all reported variant classes in a candidate region, and project selected variants onto a phylogenetic tree. Tables and figures can be downloaded, and selected analyses can be run on user-submitted gene alignments. This integrated workflow is designed for researchers studying TB evolution, drug resistance, vaccine-candidate stability, genome plasticity, and population structure in clonal bacterial pathogens.

Here we describe the construction, content, statistics, and interface of TBpop. Following a data-first structure, we first summarize the source dataset and database organization, then describe the scale and distribution of the hosted variation and population-genetic statistics, and finally present the web interface and user-analysis functions. TBpop is freely available at https://tbpop.chinacdc.cn.

## Methods

### Implementation

TBpop is a containerized web application built with a React/Vite frontend, FastAPI backend, PostgreSQL database, and background worker. Docker Compose manages separate database, API, worker, and frontend services, with runtime settings supplied through environment variables. The backend uses SQLAlchemy models and modular API routers to serve datasets. The frontend links Population, Statistics, Variation, User Analysis, and Document views, allowing users to move from population or gene-level signals to genome tracks, summary tables, downloads, and phylogeny-linked figures. Uploaded user analyses are quality-checked, queued, processed asynchronously by the worker, and returned with downloadable result artifacts.

### Data Collection

TBpop was constructed from a curated population genomic dataset of 420 clinical MTB isolates from China (**Figure 1**). In total, 3,929 MTB isolates were collected in the first national drug-resistance baseline survey, a cluster-randomized survey of tuberculosis cases in the public health system conducted between April 1 and December 31, 2007 [13]. From this survey collection, 420 representative MTB strains were selected for whole-genome sequencing based on genotype, drug-susceptibility profile, and geographic information, so that the dataset captured the diversity of epidemic strains circulating across China [15].

**Figure 1.**
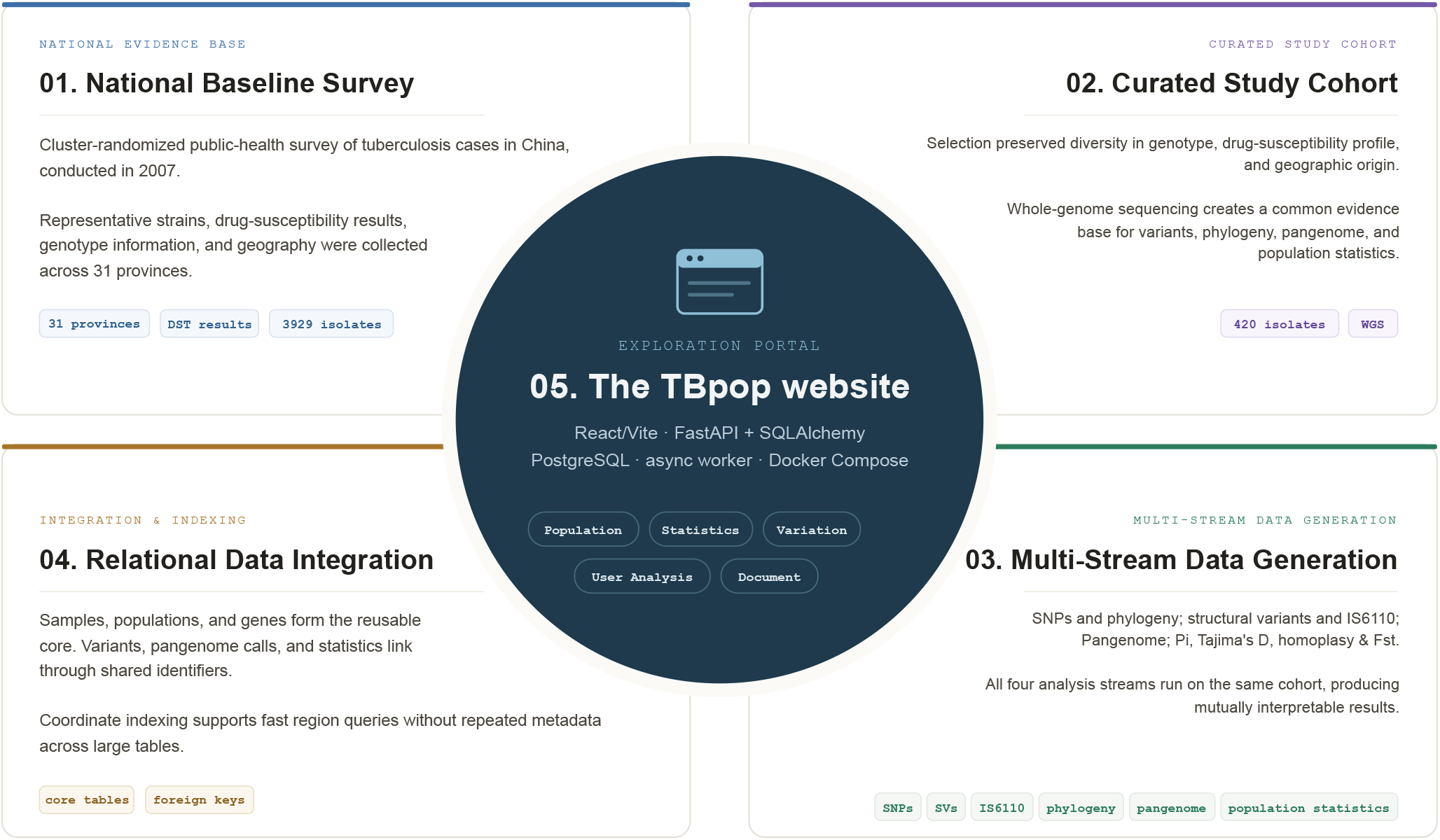
TBpop Construction Workflow. A national survey is transformed into linked genomic evidence and an exploratory web portal.

### Data Generation

All data presented on TBpop have been published in two prior studies of the 420 representative MTB strains [15,16], and detailed methodology can be found in those publications.

Briefly, short reads were mapped to the reference genome of H37Rv using BWA and SNPs were called with bcftools [17,18]. The phylogeny of the 420 strains was reconstructed from 33,220 filtered SNPs across the 420 genomes after filtering repetitive regions and low quality calls. The final tree was inferred by maximum likelihood under a GTR+G model using MEGA [19]. Genotypes of genomes were assigned based on robust SNP barcodes [20]. For population genetic statistics, SNPs were extracted from per-gene alignments derived from de novo assemblies using SPAdes and the Biopython library [21,22]. SNP functional effects and impact classes were taken directly from SnpEff annotation [23].

Eleven open source software packages were used to call insertions, deletions and inversions, including assemblytics [24], bcftools [18], breakdancer [25], delly [26], lumpy-smoove [27], minimap2 [28], svaba [29], softsv [30], tiddit [31], unimap [32], and wham [33]. After extensive filtering, only high confidence SVs were kept in the dataset. IS6110 was detected using ISdetector for both paired-end and single-end strains and only high confidence records were kept [34].

The pangenome of the 420 isolates was constructed by combining *ab initio* genome annotation using Prokka/GET_HOMOLOGUES [35,36], and blasting of annotated genes with blastn [37]. In the final pangenome, coding genes were classified into four categories: core, soft-core, shell and cloud genes. The categories describe functional gene presence rather than simple DNA detectability: core genes have valid uninterrupted open reading frames (ORFs) in all 420 clinical genomes and H37Rv; soft-core genes are interrupted or deleted in fewer than 5% of genomes; shell genes are functionally present in 3 to 400 genomes; and cloud genes are functionally present in fewer than three genomes. Thus, a gene can be physically represented in an assembly but counted as absent from the functional pangenome if it is disrupted by any type of variant such as non-sense SNPs or large deletions. If no disruptive variants were detected in a gene, but the current pipeline still failed to recover a functional gene sequence, the corresponding isolate was counted as low quality for that gene.

Tajima’s D and π were calculated per valid coding-gene alignment with DnaSP v6 [38]. π was only calculated for genes with more than one valid ORF. Tajima’s D was only calculated for genes with > 3 valid ORFs and at least one segregating nucleotide site. Tajima’s D significance is not assigned using a fixed absolute cutoff. Instead, TBpop uses the backend’s beta-distribution approximation from Tajima’s neutral-mutation test to precompute two-tailed critical values for sample-size buckets, and each gene was evaluated against the critical values corresponding to its effective sample count [9].

Homoplasy events were calculated at both SNP level and gene level using HomoplasyFinder [39]. For SNP-level homoplasy calculations, this whole-genome phylogeny provided the evolutionary scaffold: for each gene, the tree was pruned to samples with valid ORFs of this gene before repeated mutations were counted. For gene-level homoplasy calculations, the pangenome matrix presenting the presence/absence of valid ORFs was used as input.

Fst was calculated from SNP variation among lineage or subgroup assignments using HierFstat [40], with negative values set to zero and significance assessed by permutation of sample labels in the source analysis.

### Database Schema

The database is organized around a small relational core rather than storing each incoming file as an isolated dataset (**Figure 2**). Sample metadata is centered on the samples and populations tables, while reference annotation defines the genes table. Variant calls, population statistics, and pangenome categories connect back to these core records through shared integer identifiers such as sample_id, population_id, and gene_id. Coordinates are handled as a separate access dimension, allowing genomic regions to be queried by overlap rather than forcing every positional relationship into a foreign key.

**Figure 2.**
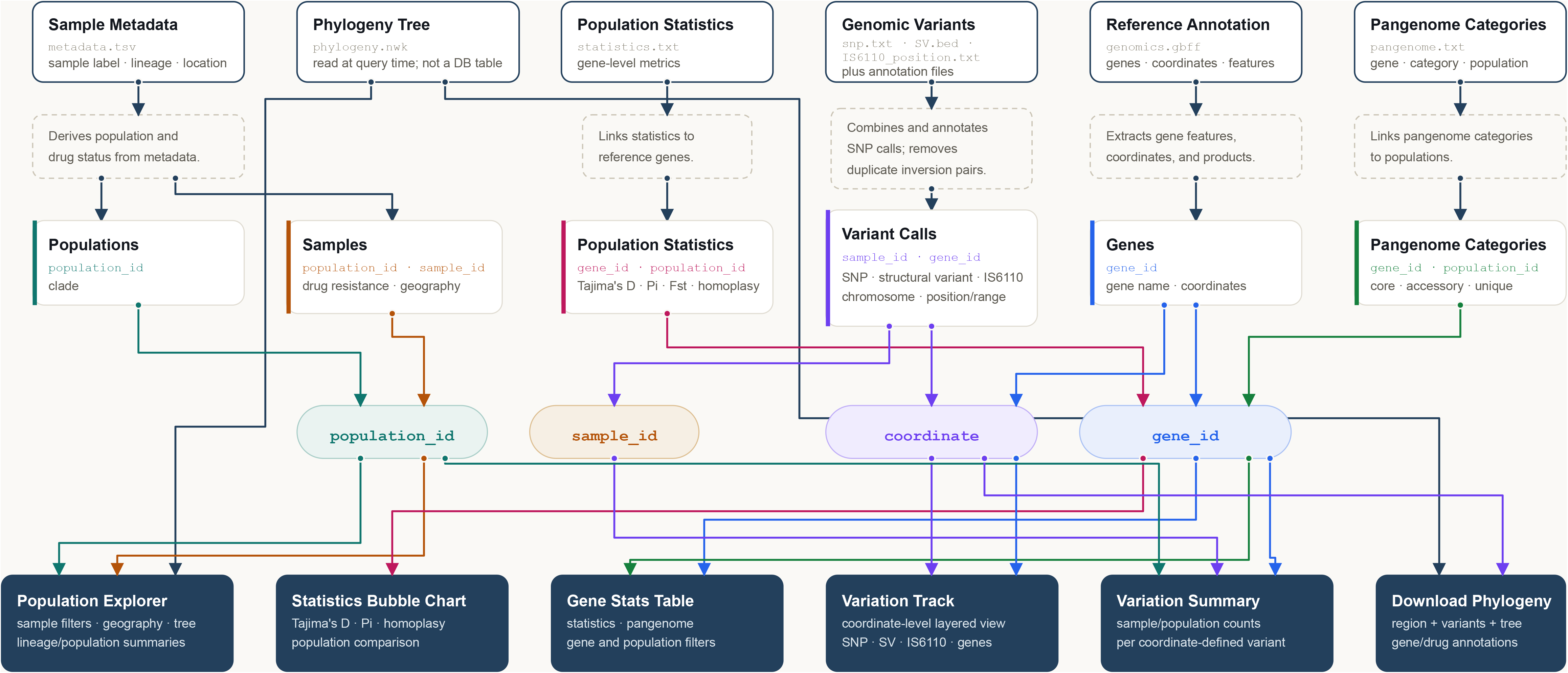
Portal Dataset Relationships. Coloured connectors trace each input dataset through its shared identifiers to the portal views that use it.

This relational design avoids duplicating sample and gene annotations across millions of variant records, keeps updates consistent, and supports focused indexes for identifier-based joins and coordinate-based region queries. Descriptive information about a sample, population, or gene is stored once in its core table, while high-volume observations—SNPs, structural variants, IS6110 positions, and gene-level statistics—reference those entities. That avoids duplicating sample and gene annotations across millions of variant records, keeps updates consistent, and supports focused indexes for identifier-based joins and coordinate-based region queries.

The design also supports the portal’s different explorer modules without creating a separate database for each view. For example, the Variation Track combines coordinate lookup with variant and gene records; Variation Summary combines variants with sample and population context; and population or gene statistics reuse the same core identifiers. The foreign key layer makes these relationships explicit and keeps the dataflow understandable even when one query spans several data types.

The schema is readily expandable when new data arrives. New samples can be added with their metadata and linked to an existing or newly created population; new variants simply reference the appropriate sample_id and, where available, gene_id. New analysis types can be introduced as additional tables that use the same identifiers, without redesigning the existing core. Likewise, new reference annotations, pangenome classifications, or summary metrics can be appended while preserving the stable sample, population, gene, and coordinate access patterns that existing portal modules depend on.

The database therefore behaves as the linked evidence model shown in **Figure 2** rather than as a set of independent tables. Population filters select samples; samples connect to per-sample SNP, SV, and IS6110 events; H37Rv coordinates connect those events to annotated genes; genes connect to pangenome category and population-genetic statistics; and the phylogeny provides the tree context for interpreting whether variants are clade-associated or repeatedly distributed. This structure supports iterative queries such as selecting a gene with significant Tajima’s D and high homoplasy, opening its H37Rv interval in the Variation Explorer, reviewing all SNP/SV/IS6110 events in that interval, and projecting selected events onto the strain phylogeny.

## Results

### Isolates Metadata

There are 420 Mycobacterium tuberculosis strains collected across 31 Chinese provincial-level regions in TBpop. The collection is dominated by lineage 2.2-modern strains (274, 65%), followed by lineage 2.2-ancient strains (70, 17%); the remainder includes lineage 4 sublineages and a small number of lineage 1 and lineage 3 strains. This composition reflects population structure of the epidemic MTB strains in China. Drug-susceptibility fields record resistance to rifampicin, isoniazid, streptomycin, ethambutol, kanamycin, and ofloxacin: 282 strains have resistance recorded for at least one drug, including 192 with both rifampicin and isoniazid resistance.

### Summary of Genetic Variants

The current variation layer contains 737,264 per-sample SNP records from reference genome alignment and *de novo* assembly alignment, representing 55,944 unique SNP positions; 93,589 per-sample SV records, representing 8,683 unique SV events; and 5,343 per-sample IS6110 insertion records, representing 957 unique insertion events.

In TBpop, approximately 51% of assembly-only SNPs (23,827/46,730) are reported as the reference genotype by bcftools. These represent sites where the de novo assembly diverges from H37Rv but bcftools finds no variant in the read-alignment data. The remaining 49% of the assembly-only SNPs occur in regions that are highly divergent from the H37Rv reference sequence where short reads are soft/hard-clipped, thus bcftools returns no genotype information for these sites. For example, a 115-bp region in *pe_pgrs17* is sufficiently divergent that reads spanning the region are soft-clipped.

Most annotated SNPs fall within coding regions and many alter protein sequences, consistent with the compact, gene-rich organization of the H37Rv reference genome (**Figure 3**). When a SNP had multiple SnpEff annotation rows because it overlapped more than one genomic feature, it was counted once using the highest-impact annotation. Under this unique-SNP counting scheme, the 56,356 annotated SNP alleles (including multi-allelic sites counted separately per alternate allele) include 681 high-impact variants, 33,005 moderate-impact variants, 16,846 low-impact variants, and 5,824 modifier variants. At the effect level, 33,005 are missense variants and 16,802 are synonymous variants; high-impact coding changes include 526 stop-gained variants, 89 start-lost variants, and 66 stop-lost/splice-region variants. These annotations allow users to separate neutral or near-neutral synonymous changes from candidate protein-altering or gene-disrupting mutations.

**Figure 3.**
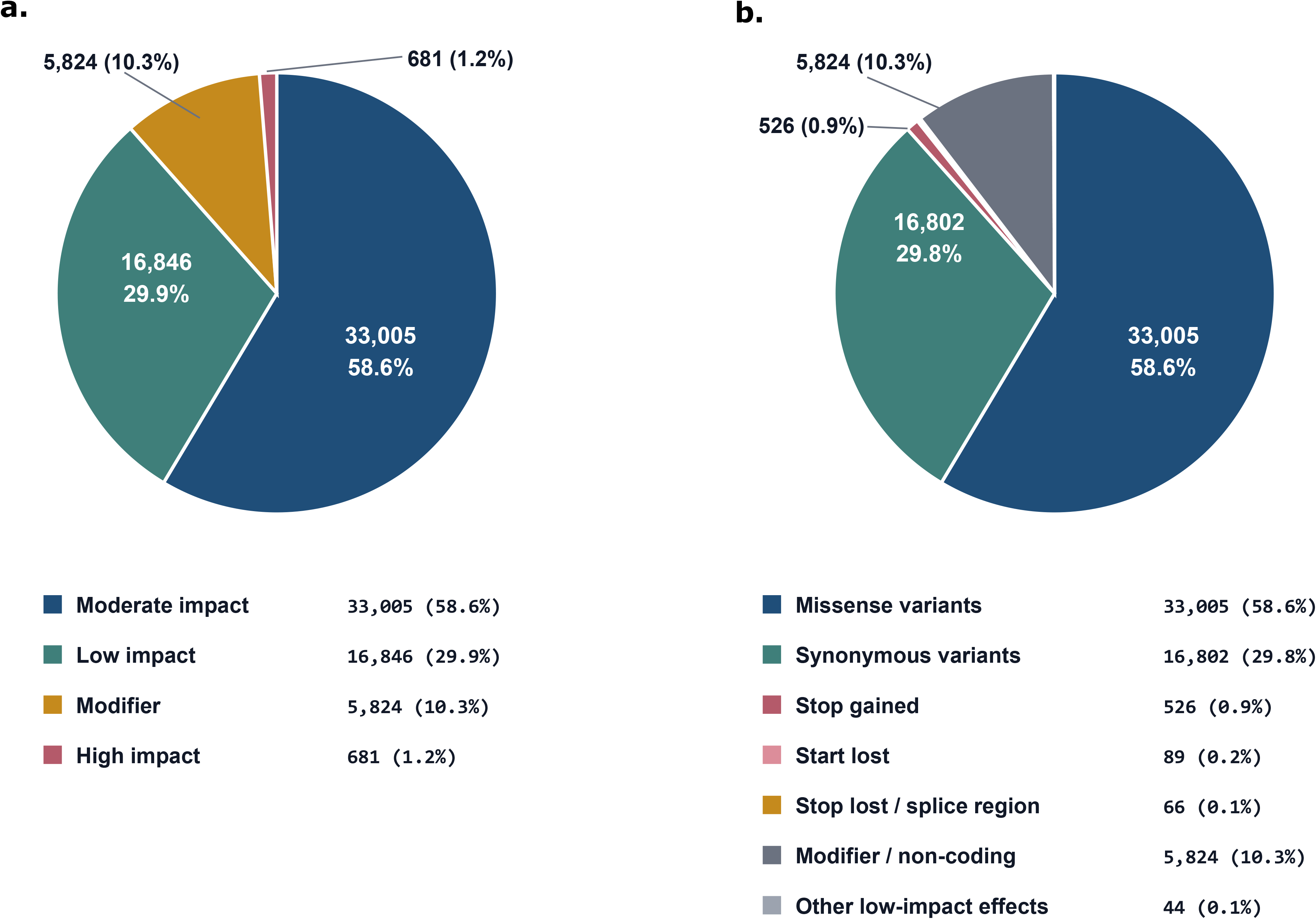
SNP annotation summary of unique alleles. **a.** impact categories of unique SNP annotations. **b.** effect categories of unique SNP annotations.

SV calls provide an additional layer of gene disruption not captured by SNPs alone. The unique SV events comprise 5,650 deletions, 2,934 insertions, and 99 inversions. At the per-sample record level, their size distribution is summarized by SV type in **Figure 4**: 52,621 records are 1-12 bp, 24,039 are 13-100 bp, 9,877 are 101-1,000 bp, 6,489 are 1,001-10,000 bp, and 563 are >10,000 bp. Large per-sample records include 148 inversions, 402 deletions, and 13 insertions above 10,000 bp. This size and type distribution helps clarify the relative contribution of different SV types across size ranges.

**Figure 4.**
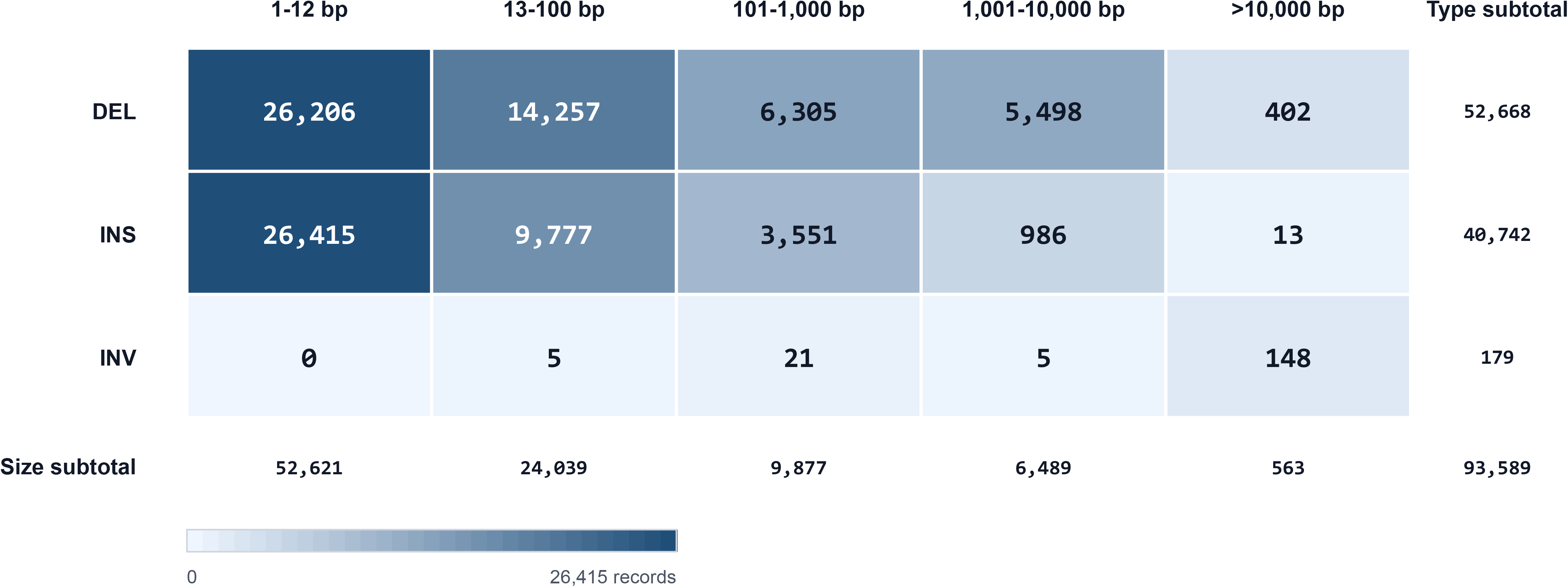
SV size distribution by event type. Cell fill uses square-root scaling; labels show exact per-sample record count.

IS6110 insertions are also frequently gene-associated. Among 890 annotated unique IS6110 insertion records, 576 overlap coding regions or genes and 314 are intergenic (**Figure 5**). The insertions are nearly balanced by orientation, with 455 forward and 435 reverse records, and most are close to full-length IS6110 elements. Because IS6110 insertions can interrupt coding sequences or alter nearby regulation, TBpop stores them as a distinct variation class rather than collapsing them into generic SV records.

**Figure 5.**
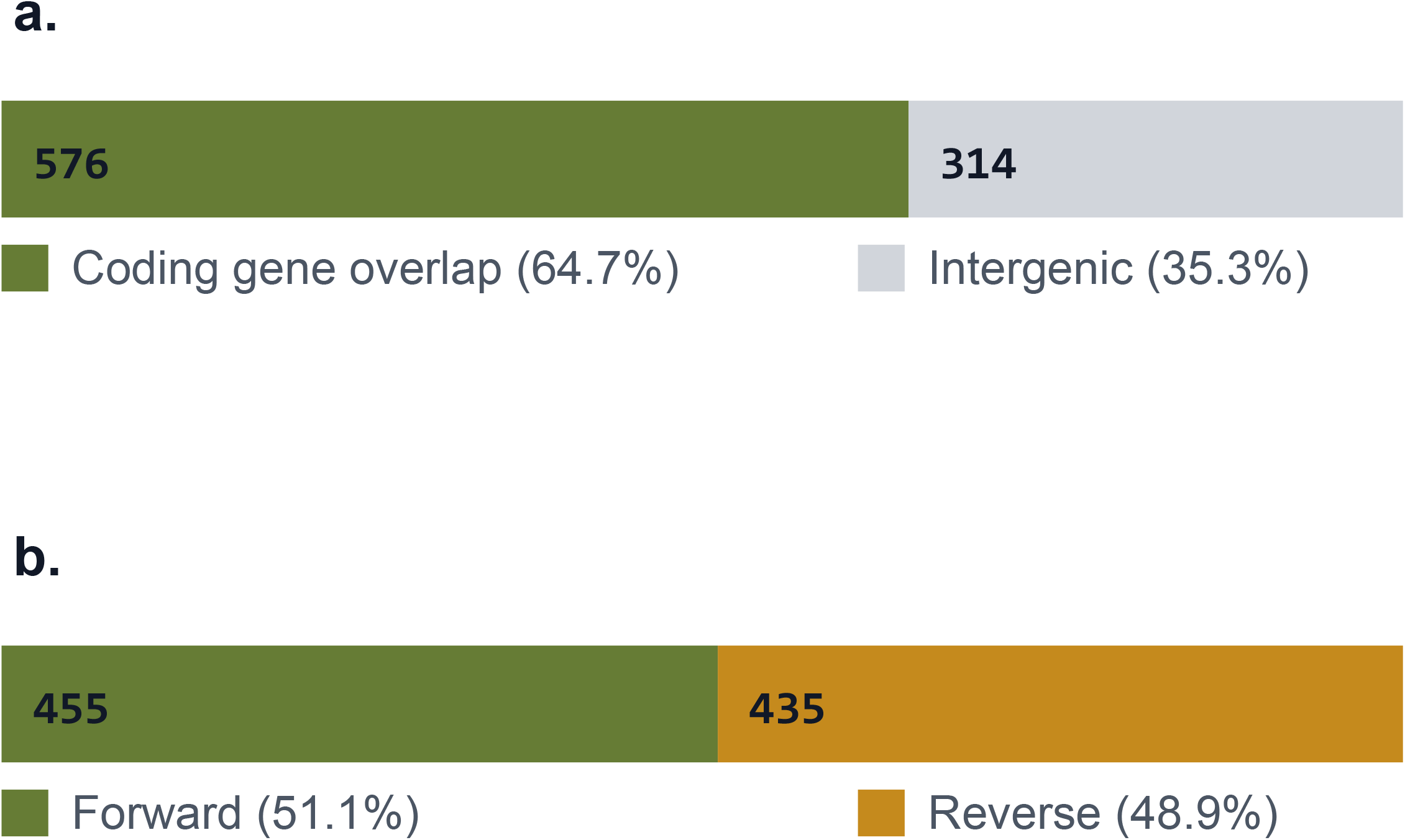
Summary of IS6110 insertions. a. annotations of IS6110 insertions in coding genes and intergenic regions. b. orientations of IS6110 insertions.

There are in total 32,379 low-quality calls in 1,328 genes. These low-quality calls are concentrated mainly in 253 genes (28,587, 88.29%) that are prone to assembly gaps when using short-read sequencing data. In the remaining low-quality calls (3,792), 3,695 (97.44%) are concentrated in 33 isolates.

### Summary of Population Genetic Statistics

Gene-level π is available for 3,889 entries (**Figure 6**). The mean π is 2.21e-4, while the median is 8.29e-5, indicating that most genes are highly conserved and that the upper tail contains a smaller set of more variable genes. In total, 768 genes have π above the genome-wide mean, 244 genes have π > 5.0e-4, 64 genes have π > 1.0e-3, and 24 genes have π > 2.0e-3. These high-diversity genes provide candidate loci for downstream inspection, especially when they also show elevated homoplasy or strong population differentiation.

**Figure 6.**
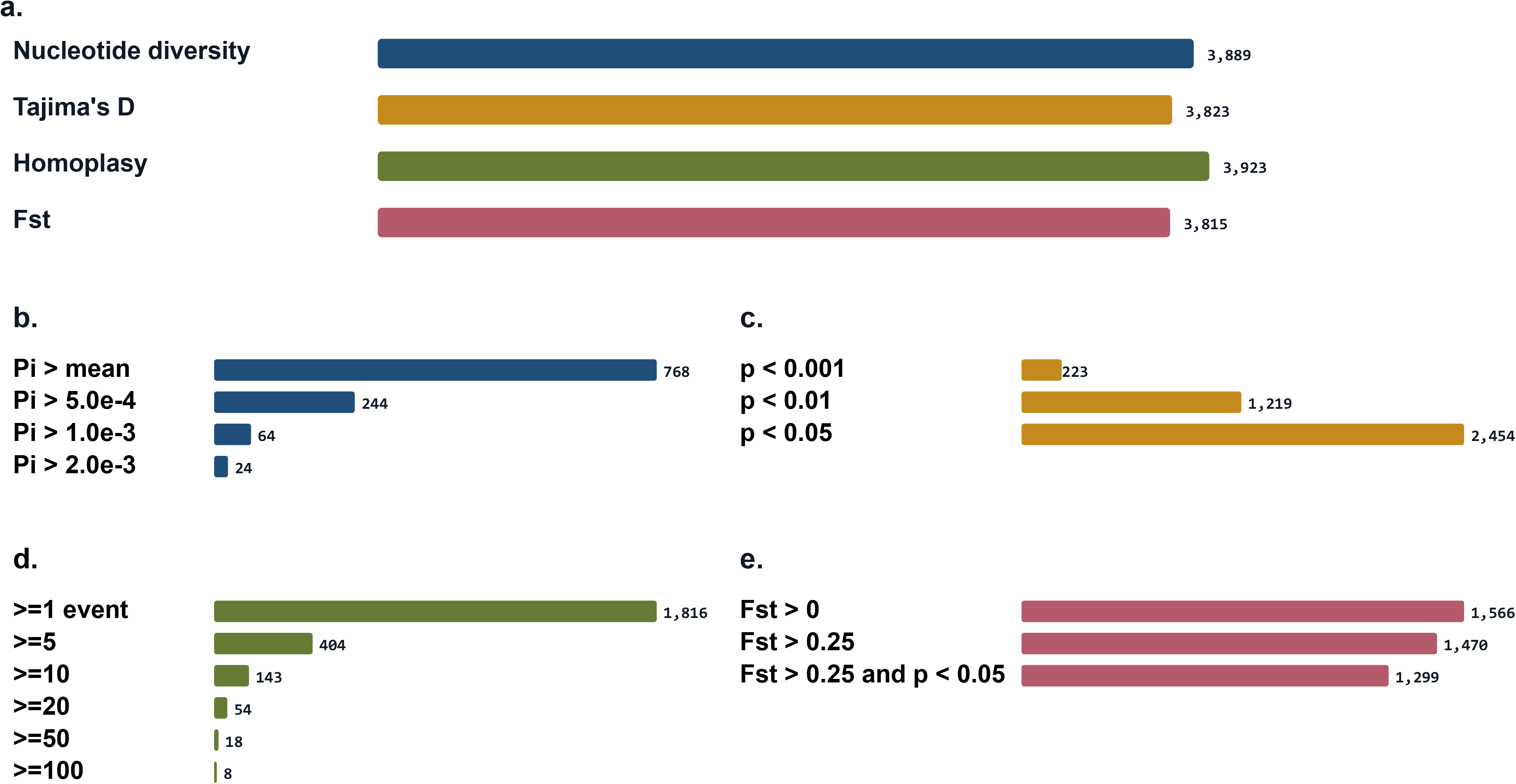
Availability and threshold counts for π, Tajima’s D, homoplasy, and Fst. **a.** availability of population genetic statistics in the 3,923 genes. **b-e.** threshold counts for the four population genetic statistics. Mean value of π is 2.2e-4 when including genes with low-quality calls. The counts are accumulative. For example, the count for genes with π > 5.0e-4 includes the genes with π > 1.0e-4.

Tajima’s D is available for 3,823 entries, fewer than the number with π. In the source pangenome study, interrupted gene copies were excluded before calculating these statistics and Tajima’s D was obtained only for coding genes with more than three valid ORFs and at least one segregating nucleotide site. In the current TBpop table, 66 entries have π but no Tajima’s D; 58 of these have zero segregating sites, and 11 have an effective sample count of three or fewer.

For the genes with Tajima’s D, values range from -2.864 to 2.069. Using the p < 0.05 thresholds, 2,454 genes fall below the lower critical value and none exceed the upper critical value; at p < 0.01, the corresponding counts are 1,219 and none. The predominance of significantly negative values is consistent with the declining population size described for this sample set, and therefore individual genes should be interpreted in combination with diversity, homoplasy, lineage structure, and variant context rather than by Tajima’s D alone.

Homoplasy counts provide a tree-aware measure of repeated mutation including SNPs and interruptive SVs. Among 3,923 entries, 1,816 genes have at least one homoplasy event (mutations that arose independently on two or more branches of the phylogeny), 404 have five or more, 143 have ten or more, 54 have twenty or more, 18 have fifty or more, and eight have at least 100. The highest homoplasy counts occur in genes including *rpoB*, *Rv0538*, *katG*, *rpsL*, *embB*, *gyrA*, *rpoC*, *Rv1769*, and *esxR*. Some of these genes are well-characterized drug-resistance gene. The remaining high-homoplasy genes are useful entry points for exploring whether repeated changes reflect drug-resistance selection, antigenic or cell-surface variation, lineage effects, mapping uncertainty, or other evolutionary processes. As noted in the Statistics Explorer gene table, genes with many unclassified or low-quality interruptions should be interpreted cautiously because missing or excluded gene copies can reduce the effective sample size and may bias homoplasy estimates for some loci.

Population differentiation is summarized by Fst for 3,815 entries. A total of 1,566 genes have Fst > 0, 1,470 have Fst > 0.25, and 1,299 have both Fst > 0.25 and Fst significance < 0.05. Genes with high Fst may contain lineage-private or lineage-enriched variation, especially when high Fst co-occurs with elevated π or homoplasy.

### Explorer Modules

TBpop is available as an open-access web portal at https://tbpop.chinacdc.cn. The interface is organized around the main analytical tasks supported by the database: browsing isolate metadata and population structure, screening genes by population-genetic statistics, inspecting genomic variation in coordinate space, generating phylogeny-linked variant figures, and running user-submitted gene analyses. The navigation bar provides direct access to the Home, Populations, Statistics, Variation, User Analysis, and Document pages (**Figure 7**).

**Figure 7.**
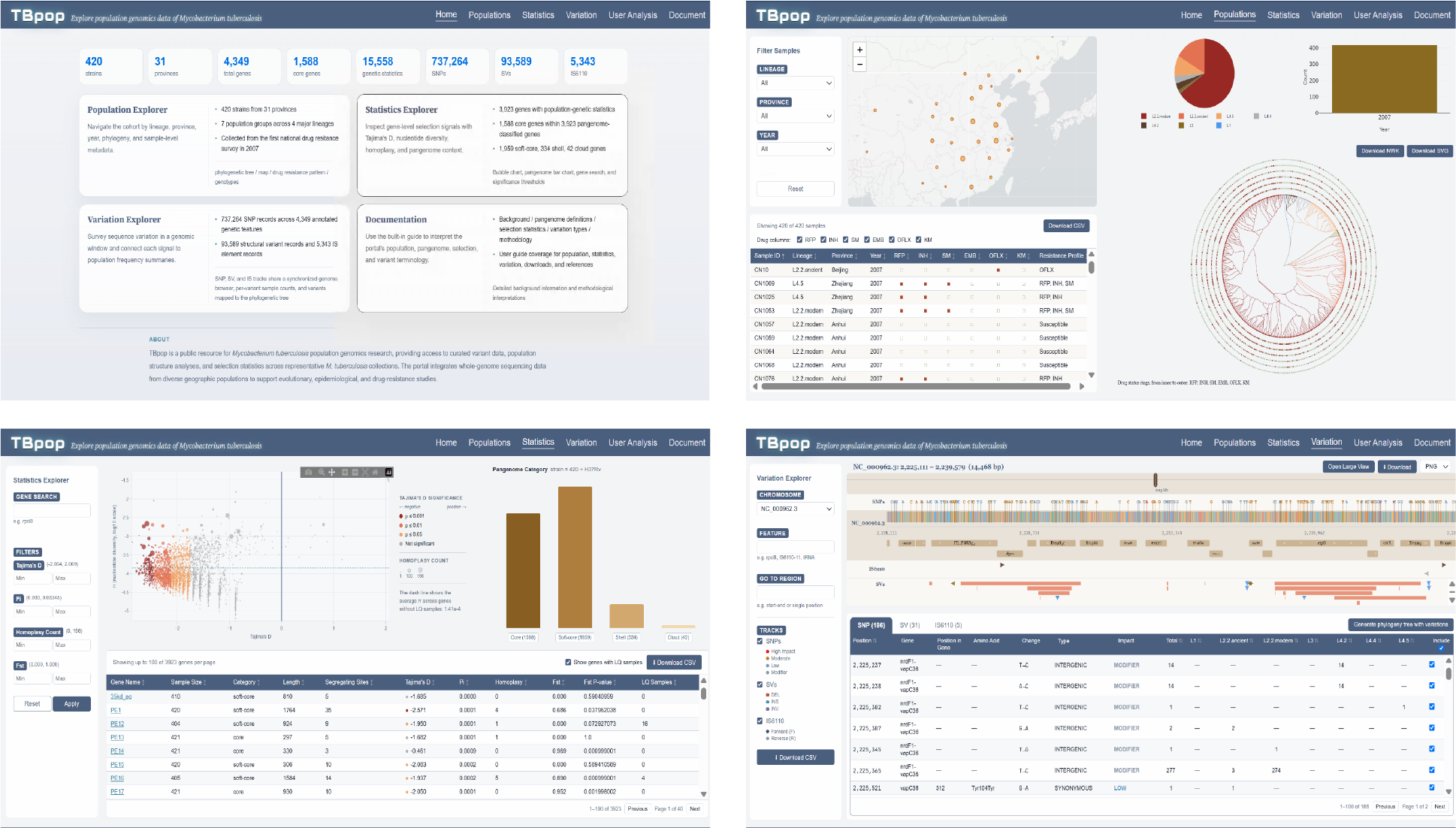
Interface of TBpop website.

The Population Explorer provides an isolate-centered view of the dataset. A geographic map, lineage summary chart, sample table, and phylogenetic tree are linked through shared filters. Users can filter isolates by lineage, province, and collection year; these filters update the map, lineage summaries, tree, and table together. This design allows users to move between geographic distribution, lineage composition, individual isolate metadata, and phylogenetic placement without switching between independent files or figures.

The Statistics Explorer provides a gene-centered entry point into the database. A bubble chart plots Tajima’s D on the x-axis and π on the y-axis, with bubble size representing homoplasy count and bubble color representing Tajima’s D significance based on gene-specific sample-size thresholds. Users can search by gene name, filter numerical ranges for Tajima’s D, π, homoplasy, and Fst, and filter by pangenome category. The accompanying gene table reports gene length, segregating sites, sample count, low-quality sample count, pangenome category, and population-genetic statistics. Left-clicking on a bubble or a gene table row highlights the corresponding entry in the other view; right-clicking navigates directly to the gene’s locus in the Variation Explorer track. The Statistics Explorer is designed to support candidate-gene discovery, for example by identifying genes with high homoplasy, high π, significant Tajima’s D, or strong population differentiation.

The Variation Explorer links gene-level signals to the underlying genomic events. SNPs, SVs, and IS6110 insertions, are displayed together in the Variation Explorer, allowing users to distinguish single-nucleotide changes from larger gene-disrupting events and insertion-sequence-mediated disruption. Users can navigate by chromosome coordinates or search directly for a gene or annotated feature. The viewer displays a configurable genomic window of up to 30,000 bp with separate tracks for sequence context, gene features, and the three types of variants. SNP, SV, and IS6110 summary tables are synchronized with the track viewer: selecting a marker highlights the corresponding table row, and selecting a row highlights the corresponding marker. This track-to-table interaction allows users to inspect all reported variants in a candidate gene or region and distinguish point mutations from larger structural or insertion-sequence events.

For phylogenetic interpretation, users can generate a downloadable phylogeny figure with selected variants mapped onto the isolate phylogeny via the Variation Explorer (**Figure 8**). Users choose which SNPs, SVs, or IS6110 insertions from the current region should be included, then generate a downloadable phylogeny-with-variation figure. This is especially useful for high-homoplasy genes because it lets users assess whether a variant is confined to one lineage, shared by a clade, or distributed across multiple branches in a pattern consistent with repeated emergence. The same region-level variant selections can also be exported as CSV files, and rendered figures can be downloaded for downstream reporting.

**Figure 8.**
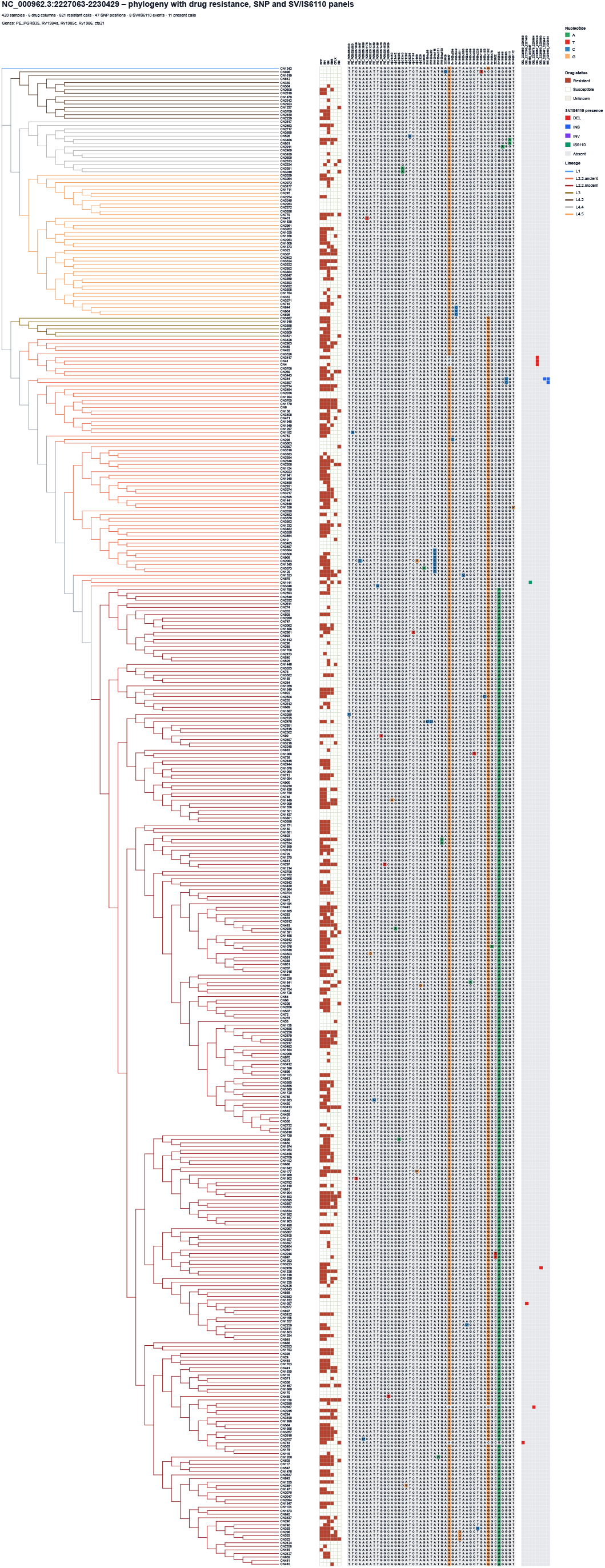
Example of downloadable phylogeny with drug-resistance and variants.

The User Analysis module extends TBpop from a static reference portal to an analysis service for user-provided genes. Users upload aligned coding-sequence FASTA files and can optionally provide a Newick tree and a genotype-definition file. Before job creation, the portal performs input checks such as duplicate-sequence detection, minimum sequence-count requirements, coding-sequence validation, tree/sample compatibility checks, and required companion-file checks for selected analyses. The workflow supports π, Tajima’s D, homoplasy, and Fst, and returns a persistent job ID for revisiting results.

User-analysis results are displayed on a dedicated result page. The page reports per-analysis job status, failed-analysis messages when applicable, creation and completion times, result expiry, statistics tables, variation summaries, per-gene tree figures, and a downloadable archive. Tree figures generated from uploaded data use the same general visual grammar as the Variation Explorer, pairing a phylogeny with gene-level SNP information so that users can interpret their own uploaded alignments in a format consistent with the reference database.

The Document page provides definitions, methodology notes, interpretation guidance, and references for the portal. It explains pangenome category definitions, SNP/SV/IS6110 interpretation, π, Tajima’s D, homoplasy, Fst, pangenome construction, variant detection, phylogenetic reconstruction, and user-analysis requirements. This documentation is integrated into the portal so that users can interpret statistics and visualization outputs without consulting separate supplementary files.

Together, these interface modules support an iterative workflow from summary statistics to mechanistic inspection. A user can identify a gene with high homoplasy and significant Tajima’s D in the Statistics Explorer, open the same gene in the Variation Explorer, inspect all SNP/SV/IS6110 events in the region, select variants of interest, and generate a phylogeny-with-variation figure to evaluate whether the events are lineage-associated or repeatedly distributed across the tree. This workflow is central to TBpop’s purpose: making complex MTB population-genomic data explorable as connected evidence rather than as isolated tables.

### Case Study: ESX System in MTB

Statistical filtering in TBpop enables systematic candidate gene selection. For example, in TBpop, 13 genes have high homoplasy counts (>=50), π above the genome-wide mean, and Tajima’s D below the gene-specific p < 0.05 lower critical value, including rpoB, Rv0538, katG, and embB. A user can start from this statistical pattern, select a candidate gene, inspect all SNP/SV/IS6110 events in the corresponding region, and project selected variants onto the phylogeny to evaluate whether variants are lineage-clustered or repeatedly distributed across independent branches.

An interesting case study is the ESX system in MTB. Twenty-three ESX genes are annotated in the H37Rv genome, and they show striking evolutionary divergence:

- High-homoplasy ESX genes (n=7): *esxR*, *esxI*, *esxS*, *esxP*, *esxO*, and *esxW* show robust signals of positive selection (homoplasy ≥9), with *esxN* showing intermediate homoplasy (n=8). These genes are all shell genes and are non-functional in varying numbers of isolates. These genes are likely to experience adaptive pressure related to host-pathogen interactions or inter-strain competition.
- Highly conserved ESX genes (n=10): Ten ESX genes, including *esxA* and *esxB*, harbor zero homoplasy events despite sampling 420 globally distributed strains. Except three with high number of low-quality samples, all are core or soft-core genes present across the entire dataset. This extreme conservation suggests these genes encode essential functions that tolerate little sequence variation, likely involving critical virulence mechanisms or cell survival pathways.
- Intermediate group (n=6): The remaining six ESX genes show low homoplasy levels (2–4 events), occupying a spectrum between conservation and adaptation.

This bifurcation reflects distinct evolutionary roles within the ESX system and have key implications in the development of new vaccine and new diagnostic tools: conserved ESX genes are strong vaccine candidates and potential targets for broadly effective interventions; divergent ESX genes may serve as lineage markers or drivers of phenotypic diversity; adaptive genes are biomarkers for lineage identification or drivers of phenotypic diversity. Especially, *esxR* is among the most divergent ESX genes, featuring 22 unique SV and IS6110 insertion events across 162 of the 420 strains, but only two SNP loci. This dissociation between SV-dominated and SNP-dominated evolution within a single gene is not universal and highlights the importance of integrated variant analysis.

## Discussions

TBpop is an open-access population genomics portal that integrates isolate metadata, pangenome categories, SNPs, structural variants, IS6110 insertions, phylogeny, and gene-level statistics for 420 clinical *Mycobacterium tuberculosis* isolates from China into a single interactive resource. Its core utility lies in linking complementary evidence types — users can move from population-genetic signals such as elevated homoplasy or significant Tajima’s D directly to the underlying SNP, SV, and IS6110 events, and visualize selected variants on the isolate phylogeny without reconciling separate files or tools. This integrated workflow is particularly suited to studying a clonal pathogen where evolutionary interpretation requires combining gene-level summaries with genomic context and phylogenetic structure.

TBpop is intended to complement, rather than replace, existing TB resources. Resistance-focused databases and general MTB variant catalogues remain essential for interpreting known antimicrobial-resistance mutations and accessing large-scale sequence collections. TBpop addresses a different need by emphasizing integrated population-genomic interpretation, structural variation, IS6110-mediated genome disruption, pangenome status, and exploratory statistics in a single web environment. The User Analysis module further extends the portal beyond the hosted reference dataset by allowing users to analyse their own coding-sequence alignments and compare results using the same general framework.

Several developments are planned for future versions. We aim to expand the curated dataset with additional representative MTB genomes from national surveillance and collaborative studies, while retaining the current emphasis on high-quality metadata, transparent provenance, and reproducible analysis. Broader sampling will make it possible to compare population-genetic patterns across regions, time periods, lineages, and resistance backgrounds. We will also improve the representation of non-SNP variation, including structural variants and mobile-element insertions, because these variation classes remain underrepresented in many web-accessible MTB resources. Additionally, we plan to add richer comparative functions, such as cross-dataset gene comparison, lineage-specific summaries, and expanded export options for downstream analysis. Long-term maintenance will focus on usability, interoperability, and reproducibility. The user-analysis workflow will also be refined to support diversified formats of input files and additional visual outputs where appropriate. These developments are intended to make TBpop useful not only as a browser for one dataset, but also as a durable framework for comparative MTB population genomics.

In summary, TBpop makes MTB population-genomic data explorable as linked evidence rather than as isolated result files. Its combination of curated clinical isolates, multiple variant classes, pangenome information, population-genetic statistics, phylogenetic visualization, and user-submitted analyses provides a practical resource for researchers studying TB evolution, drug resistance, genome plasticity, and candidate gene stability. TBpop is freely available at https://tbpop.chinacdc.cn and will be updated as new datasets, analysis methods, and community needs emerge.

## Data Access

All whole genome sequencing data used in this project can be accessed from NCBI database via accession number PRJNA573798.

## Ethics Statement

This project does not involve human participants or the use of personal information and is based on previously published studies which have been approved by the Ethics Review Committee of the Chinese Center for Disease Control and Prevention.

